# *Brd4::Nutm1* fusion gene initiates NUT carcinoma *in vivo*

**DOI:** 10.1101/2023.07.29.551125

**Authors:** Dejin Zheng, Ahmed Elnegiry, Chenxiang Luo, Mohammed Amine Bendahou, Liangqi Xie, Diana Bell, Yoko Takahashi, Ehab Hanna, George I. Mias, Mayra F. Tsoi, Bin Gu

## Abstract

Nut carcinoma (NC) is an aggressive cancer with no effective treatment. The majority (70%) of NUT carcinoma is associated with chromosome translocation events that lead to the formation of a *BRD4::NUTM1* fusion gene. However, because the *BRD4::NUTM1* gene is unequivocally cytotoxic when ectopically expressed in cell lines, questions remain on whether the fusion gene can initiate NC. Here, we report the first genetically engineered mouse model (GEMM) for NUT carcinoma recapitulating the human mutation. By stochastically inducing a chromosome translocation mirroring the human event, we demonstrated that the *Brd4::Nutm1* fusion gene could induce aggressive carcinomas in mice. The tumors present histopathological and molecular features similar to human NC, with an enrichment of undifferentiated cells. Similar to the reports of human NC incidence, *Brd4::Nutm1* can induce NC from a broad range of tissues, demonstrating that its oncogenic potential is not lineage-restricted. The consistent induction of tumors of squamous phenotypes, even from ductal epithelial and mesenchymal tissues, demonstrated a strong reprogramming activity of BRD4::NUTM1. The new mouse model provided a critical preclinical model for NC and opens new opportunities for understanding the oncogenic mechanism and developing new therapies.

## 1. Introduction

NUT carcinomas (NC), formally named NUT Midline Carcinomas (NMC), are poorly-differentiated squamous cell carcinomas that often arise within midline organs^1–3^. Other primary sites, including skin^4^, pancreas^5^, pelvic organs^6,7^, and soft tissues^8,9^, have also been reported. They are aggressive tumors that typically present with metastatic disease at diagnosis^10^. The prognosis of NC is dismal, with an overall survival time of 6.5 months, even with intensive treatment, including surgery, radiation, and chemotherapy^10^. NC is rare; a recent estimation suggests that NC accounts for approximately 0.21% of all cancers, with an estimated 3500 new cases emerging in the U.S. each year^11^. However, because NC lacks a distinctive diagnostic feature and is not widely recognized by many clinicians around the world, the case rates are likely underestimated. NC is more prevalent in young people, with a median incidence age in the 20s; it causes a dramatic loss of life^12^. Thus, it is critical to understand the pathogenesis of NC, which will lead to better treatment for NC patients.

Clinical studies have established a strong association of NC with chromosome translocation events that produce fusion genes involving a testis-specific gene known as Nuclear protein in testis (NUTM1)^13^, with various epigenetic factors, including BRD4 (70% of cases), and BRD3, NSD3, ZNF532, ^1,4,9,14–17^. Limited genomic sequencing and karyotyping analysis suggested that these fusion genes, likely acting as an abnormal epigenetic modifier, can drive NC without other mutations in a quiet genomic landscape^18^. At the molecular level, it was suggested that, by recruiting the P300 acetyltransferase (HAT) to ectopic sites, BRD4::NUTM1, and the other fusion proteins, drive malignant transformation by activating oncogenes and epithelial progenitor genes such as *cMYC*, *SOX2* and *TP63*^19–23^.

However, when ectopically expressed in non-NC cell lines, *BRD4::NUTM1* is unequivocally cytotoxic^2,24,25^, contradictory to the proposed oncogenic functions. To reconcile these contradictory observations, we consider two possible explanations. First, many fusion oncogenes have been shown to require a specific cell lineage status to achieve cell transformation and are otherwise toxic to other cell types. This is coined as a concept of ‘cellular pliancy’ first in pediatric cancer and then in cancer in general^25–27^. Second, it could simply be that the over-expression system used on cell lines expresses the *BRD4::NUTM1* oncogene at a too-high level. It is not uncommon to observe gene dosage-dependent toxicity of oncogenes^28,29^. Thus, the discrepancy in expression levels could be a simple explanation for the cytotoxicity of the *BRD4::NUTM1* oncogene.

To recapitulate the NC oncogenesis in an experimental model in which both critical parameters are controlled, we developed a genetically engineered mouse model (GEMM). Using cre-lox-mediated chromosome translocation, the model ensured the recapitulation of human-mimicking chromosome translocation and established syntenic control of the resultant *Brd4-Nutm1* fusion gene. Taking advantage of the intrinsic low frequency of cre-lox mediated interchromosome translocation^30–33^, the model recapitulated the stochastic nature of fusion gene formation. Using various cre-drivers, we demonstrated a broad oncogenic activity of BRD4-NUTM1 across tissues, covering all reported sites of human NC. BRD4-NUTM1 drives aggressive squamous cell carcinoma with various degrees of differentiation, consistent with human NCs. Further, the model demonstrated a strong intrinsic metastasis potential of NC, recapitulating a key feature of human NC. Thus, we reported the first GEMM that recapitulates the intrinsic genetic lesion and oncogenic process of human NC. The model will create tremendous opportunities for understanding and developing treatments for this devastating disease.

## 2. Results

### 2.1 *BRD4::NUTM1* induces NC-like tumors from epithelial progenitor cells

We designed and developed a mouse model to model the NC initiation event using 2C-HR-CRISPR^34^ (Figure 1A). Loxp sites were inserted into the appropriate introns of the mouse *Brd4* and *Nutm1* genes to allow stochastic induction of a reciprocal t(2;17) chromosome translocation forming the Brd4-Nutm1 fusion gene by Cre recombinase. To track the development of NC *in vivo*, we inserted a Luc2TdTomato reporter cassette downstream of the endogenous *Nutm1* coding sequence, separated by a T2A self- cleaving sequence^35^. The Luc2TdTomato reporter expressed specifically in *Nutm1*- expressing post-meiotic spermatids as expected (Supplementary Figure 1 A and B). For brevity, mice carrying both loxp sites and the reporter cassette linked to *Nutm1^loxp^* are designated NC translocators (NCT). From this point, unless otherwise explained, NCT^+/wt^ refers to *Brd4*^loxp/wt^; *Nutm1* ^loxp/wt^ whereas NCT^+/+^ refers to *Brd4*^loxp/loxp^;*Nutm1*^loxp/loxp.^

**Figure 1:**
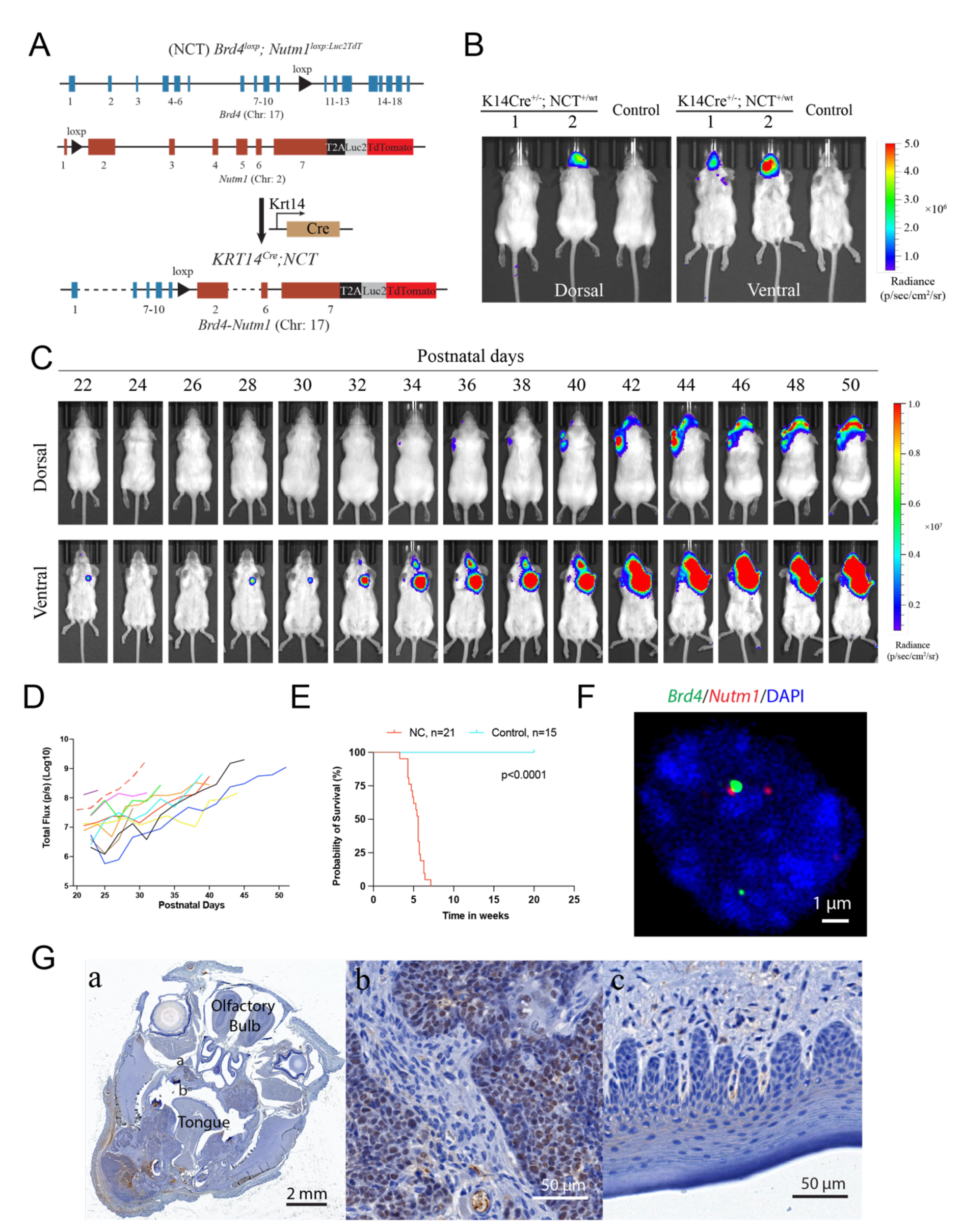
Generation of the NC GEMM: A. Genetic design of the Nut carcinoma translocator (NCT) and the generation of the Krt14Cre induced mNC. B. Representative BLI images of mice with head NC. C. BLI images of the growth kinetics of mNC. D. Growth curves of mNC in 8 mice quantified by BLI signals; each curve represents one mouse. E. Kaplan Meier survival curve of Krt14Cre induced mNC mice. P value calculated by log-rank test. F. DC-FISH of the *Brd4* (Green) and *Nutm1* (Red) on mNC. G. A representative NUTM1 IHC image of NC bearing whole mouse head (a) and zoom-ins on the tumor (b) and normal oral mucosa (c) regions.

Because around 40% of human NCs are poorly differentiated, squamous cell carcinoma (SCC) arose in the head-and-neck region and expressing markers reminiscent of basal progenitor cells^3,36–38^, we first tested NC induction using the *Krt14Cre* allele that expresses the Cre recombinase in ectoderm-derived basal progenitor cells^39^ (supplementary Figure 1C). At weaning age, KRT14Cre^+/-^; NCT^+/wt^ mice developed BLI-positive head tumors at 100% penetrance (Figure 1B and Supplementary Figure 1D)(n>50 mice), with some mice also developing skin tumors. These tumors grew exponentially (Figure 1D) and rapidly led to a moribund state (followed by humane euthanasia) (Fig. 1E), consistent with the aggressive nature of human NC, likely due to the large tumor burden in the oral cavity causing obstruction of the upper airway and digestive tracts. Upon autopsy, tumors were identified in the oral cavity (mandible, maxilla, and tongue), salivary glands, eyelids, ear canal, and skin (Supplementary Figure 1E); all of these locations have been reported in human patients and consistent with the expression pattern of Krt14Cre.

We validated the formation of the t(2;17) chromosome translocation by 1)duo color fluorescent in situ hybridization (DC-FISH) (Figure 1F); 2) Sanger sequencing of junction sequence (Supplementary Figure 1F); 3) karyotyping (Supplementary Figure 1G); and 4) fusion calling in whole genome sequencing (Supplementary figure 1H).

Furthermore, immunohistochemistry (IHC) analysis against NUTM1 detected pervasive nuclear expression of the BRD4-NUTM1 protein in the tumors (Figure 1G and Supplementary Figure 1I). No BRD4-NUTM1 was expressed in NC surrounding normal epithelial or tumor-associated stromal cells (Figure 1G), confirming the low frequency and stochastic nature of the chromosome translocation. The above-described data suggest that the t(2;17) chromosome translocation that creates *Brd4::Nutm1* can induce aggressive tumors from mouse epithelial basal cells. Because the mouse tumors satisfy two key diagnostic criteria applied to humans – the detection of the chromosome translocation by FISH and the detection of BRD4::NUTM1 proteins by IHC^15,40^, we call these tumors mouse NUT Carcinoma (mNC).

### 2.2 Mouse NCs Present Histopathological and Molecular Features of Human NC

Most human NCs are categorized as poorly differentiated squamous cell carcinomas, characterized by primitive round cells with variable areas of abrupt squamous differentiation and keratinization^11^. This heterogeneous phenotype has caused profound confusion in diagnosing NC. Also, the intra and inter-tumor heterogeneity can create a plasticity space for therapy resistance. As shown in Figure 2A, mNCs consist of large areas of basaloid, polygonal, or spindle-shaped cells. Many tumors exhibit areas of squamous cell differentiation and keratinization that occupy variable proportions of the tumor. These features are consistent with the histopathological observations in the clinical setting. Furthermore, a board-certified veterinary pathologist scored 50 cases of mouse NC arising in the head-and-neck and skin sites and concluded that the mouse NCs present a wide range of degrees of squamous cell differentiation (Figure 2B), recapitulating the human pattern. Notably, even within a single mouse, individual tumors can display variable morphology (Figure 2C), demonstrating intra and inter- tumor heterogeneity within the same genetic background. Thus, mNCs reproduced the morphological heterogeneity of human NC.

**Figure 2:**
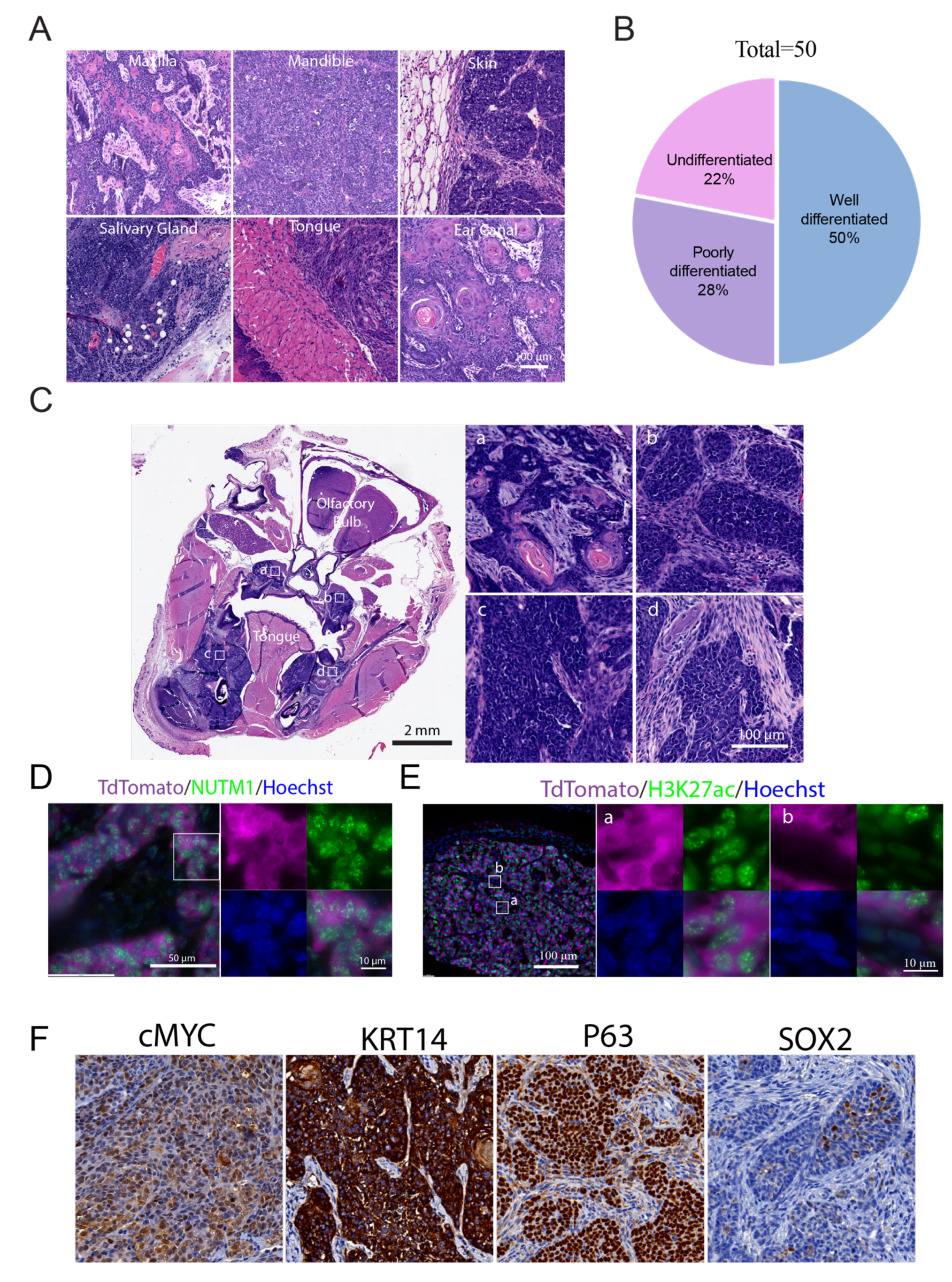
Histological and molecular characteristics of mNCs: A. Representative H&E staining images of Krt14Cre mNC tumors at different anatomical locations. B. The distribution of pathological features of mNCs. C. Tumors formed in the same mouse presenting variable histological features. D. Representative image of IF staining of NUTM1 in mNC showing nuclear foci. TdTomato staining demarcates tumor cells E. Representative images of IF analysis of H3K27ace in mNCs, depicting nuclear foci in tumor cells (b) versus a diffused nuclear stain in peri-tumor stromal cells (c). F. Representative IHC images of cMYC, KRT14, P63, and SOX2 in oral mNCs.

A few key molecular features define human NC. First, the BRD4-NUTM1 proteins form nuclear foci, possibly as biological condensates mediated by the highly disordered sequences in NUTM1, in human NC^41,42^. We detected such foci in mNCs through both Immunofluorescence (IF) and IHC (Figure 2D and Supplementary Figure 2A). Second, human NC tissues and cell lines demonstrated nuclear foci of Histone 3 lysine 27 Acetylation (H3K27ace), likely caused by the recruitment of P300 Histone Acetyltransferase (HAT) by NUTM1^19^. We detected such foci in mNCs through IF (Figure 2E and Supplementary Figure 2B). Third, human NCs express other hallmark genes, including the basal progenitor cell markers P63, Krt14, SOX2, and oncogene cMYC. These genes have also been implicated as required for the progression of NC^20,22,23^. The majority of oral mNC cells express P63, Krt14, and cMYC (Figure 2F, Supplementary Figure 2C). However, SOX2 is expressed in a small proportion of NC cells with a clustered pattern in all tested mouse NCs (Figure 2F and Supplementary Figure 2C). In clinical pathology practice, SOX2 is considered a stem cell marker in NC and is expressed in various proportions of NC cells^23,43^ , consistent with the mouse data. Whether the SOX2-positive NC cells represent a cancer stem cell population is a critical question to be investigated in the future.

### 2.3 mNCs are aggressive poorly differentiation cancer at the transcriptomics level

To gain a global understanding of how transcriptomics reprogramming promoted aggressive mNC, we performed bulk RNA sequencing analysis on six normal oral mucosa (control) and six oral mNC (tumor) samples. As shown in Supplementary Figure 3 A-C, the RNA sequencing data is of high quality, with a clear separation of the six normal oral mucosa and the six mouse NC repeats. The analysis revealed 3065 upregulated genes and 2654 downregulated genes in mNCs, demonstrating pervasive transcriptomics reprogramming (Figure 3A). Gene set Enrichment Analysis (GSEA) against the Hallmark database revealed the activation of many pathways that may contribute to an aggressive phenotype, including cell proliferation and mitosis, DNA repair, MYC target pathways, and metabolic reprogramming (Figure 3B). Notably, downregulated genes are specifically enriched for immuno-stimulatory pathways (Figure 3B).

**Figure 3:**
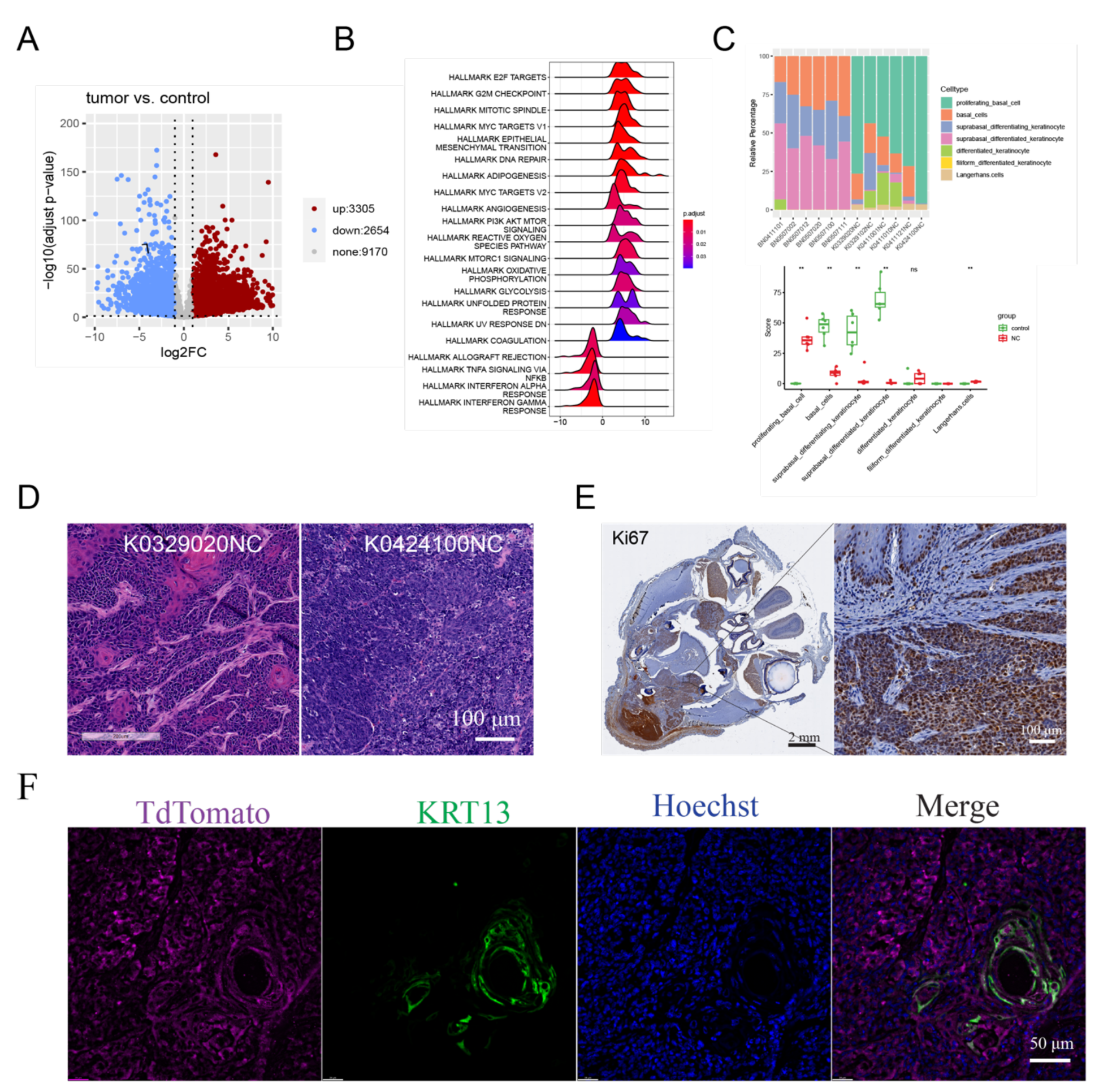
mNCs largely consist of proliferative progenitor cells. A. Volcano plot of differentially expressed genes in mNCs compared to normal oral mucosa. B. GSEA enrichment of HALLMARK pathways in mNCs. C. CIBERSORT deconvolution of the transcriptome of mNCs according to the *Tabula Muris* dataset. D. Representative histology images of mNCs that enriches the proliferative basal cell profile at different level. E. A representative Ki67 IHC image of mNCs. F. A representative IF image of NC cells (tdTomato) and the ones expressing keratinization marker KRT13.

To understand whether the poorly differentiated morphological traits are represented at the transcriptomics level, we used CIBERSORT to deconvolute the bulk RNA sequencing data using published single-cell RNA sequencing (scRNA-seq) signatures^44,45^. As reference data, we used the published scRNA-seq data on the oral mucosa epithelial from the *Tabula Muris* project^46^, which represents the cells from which the NC tumors are derived. As shown in Figure 3C, compared to the normal oral mucosa samples, there is a strong enrichment of the proliferating basal cell expression profile in the NC samples. CIBERSEQ analysis using another independently produced mouse oral mucosa scRNA-seq reference data produced similar result^47^ (Supplementary Figure 3D). The CIBERSORT results are consistent with the histological features of these tumors; for example, case K0329020 showed more differentiated histology, while K0424100NC is more undifferentiated (Figure 3D). At the molecular level, the overwhelmingly enriched proliferating basal cell profile is consistent with the pervasive expression of basal cell markers (Figure 2). Consistent with its highly proliferative nature, mNC cells overwhelmingly expressed the cell proliferation marker Ki67 (Figure 3E). In addition, the differentiated keratinocyte marker KRT13 was only detected in small regions with apparent keratinization (Figure 3F). Thus, the *Brd4::Nutm1* expression trapped most NC cells in a proliferative progenitor status, likely contributing to its aggressive nature.

### 3.4. *BRD4::NUTM1* can induce aggressive malignancy from a broad range of tissues

In humans, although most NCs were detected within the head and neck or thoracic tissues^3^, there were reports of carcinoma carrying the *BRD3/4-NUTM1* fusion genes in many other epithelial tissues, such as the pancreas^5^, the kidney^48^, and the pelvic organs^7^; and even mesenchymal tissues as undifferentiated soft tissue tumors^8^. This broad tissue distribution is a unique property of NC compared to many other fusion gene-driven cancers and indicates that the *BRD4::NUTM1* fusion gene may transform a broad range of cell types covering all three germ layers. However, due to the limited number of cases, it is unclear whether the reported *NUTM1* fusion genes are the cancer driver or background mutations irrelevant to oncogenesis in these cases. We tested it using the GEMM.

There are two cases of NC involving the pancreas reported in very young children^5^. The congenital nature of these cases suggests that an embryonic progenitor was transformed. Thus, we crossed the Pdx1Cre allele^49^, which drives recombination in endoderm-derived embryonic ductal progenitor cells into the NCT mouse line.

Pdx1Cre;NCT^+/-^ mice developed pancreatic tumors with 100% penetrance(18/18) (Figure 4A, B and supplementary Figure 4A). The pancreatic mNCs are fast-growing SCC (Supplementary Figure 4B) presenting all histopathological and molecular markers as oral mNCs (Figure 4C and Supplementary Figure 4B). Beyond the epithelial tissue, a few cases of soft tissue tumors have been reported to carry BRD3/4-NUTM1 fusion genes, suggesting that the fusion gene can also collaborate with mesoderm-derived mesenchymal gene regulatory networks (GRNs) to drive NC^8^. To test this, we used a Prrx1Cre to drive the chromosome translocation in the fetal limb mesenchyme cells^50^.

**Figure 4:**
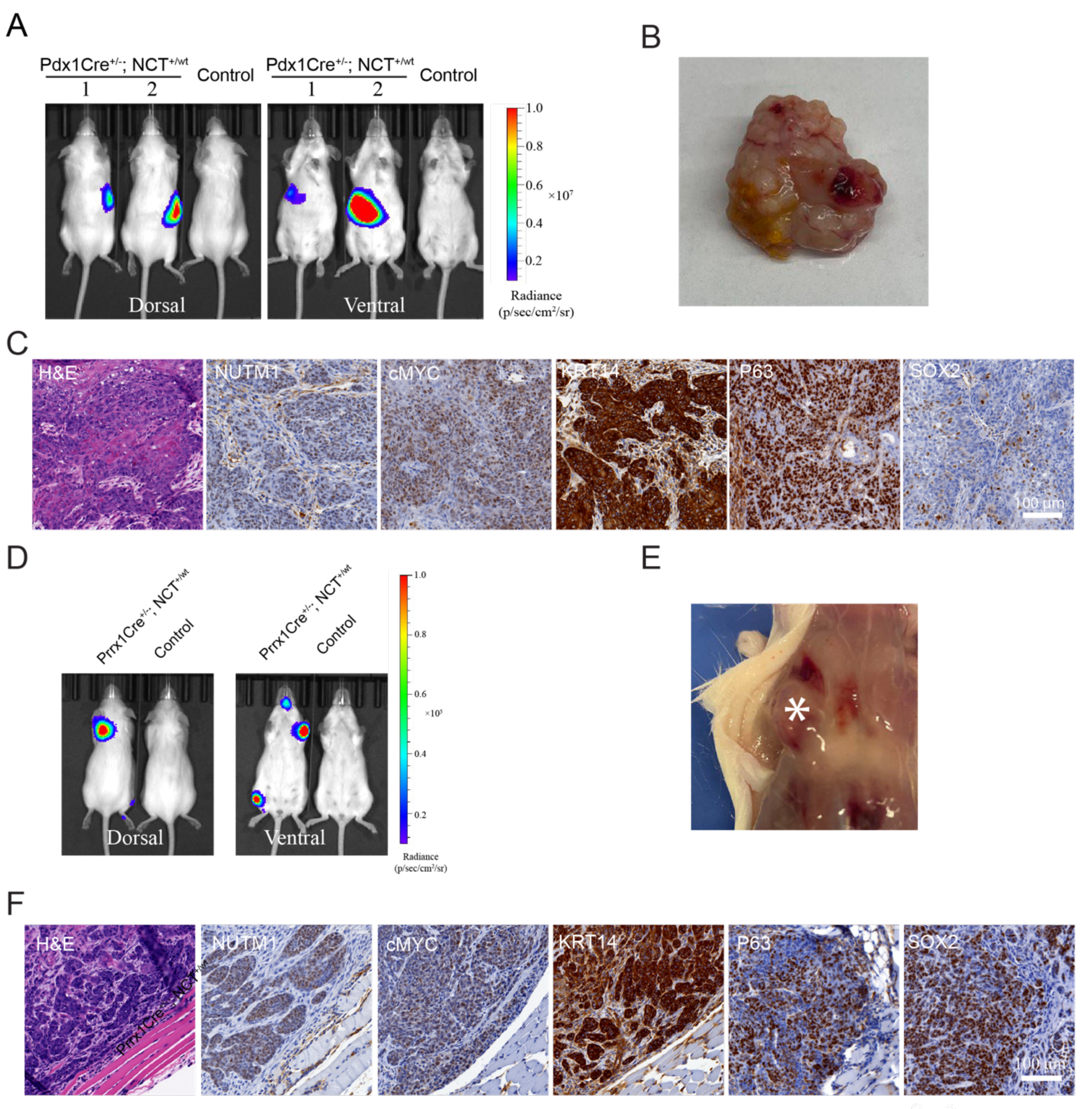
*Brd4::Nutm1* initiates pancreatic and soft tissue tumors. A. Representative BLI images of mice with pancreatic mNCs. B. A representative macroscopic image of pancreatic mNCs. C. Representative H&E, and IHC images of NUTM1, cMYC, KRT14, P63, and SOX2 in pancreatic mNCs. D. Representative BLI images of mice with soft tissue mNCs. E. A representative macroscopic image of soft tissue mNCs. F. Representative H&E, and IHC images of NUTM1, cMYC, KRT14, P63, and SOX2 in pancreatic mNCs.

These mice developed fast-growing tumors within limbs at 100% penetrance (9/9) (Figure 4D, E and Supplementary Figure 4C). The tumors are also fast-growing SCC presenting all histopathological and molecular markers of NC (Figure 4F and Supplementary Figure 4D). The ability to induce SCC from pancreatic ductal progenitors that normally initiate adenocarcinoma and mesenchymal progenitors that normally initiate sarcomas demonstrated a strong reprogramming activity of BRD4::NUTM1^49,51^.

The clonal and stochastic nature of the translocation induction provided a unique advantage of modeling the reality of stochastic gaining of the fusion gene across tissues in humans. We crossed the NCT mouse line with the NLS-Cre mouse line that constitutively expresses Cre in all mouse tissues^52^. This cross will induce *BRD4::NUTM1,* forming chromosome translocation stochastically across tissues. The NLS-Cre; NCT^+/-^ mice developed tumors detected by BLI signals at high penetrance (24/25) (Supplementary Figure 5A). Autopsy guided by BLI imaging identified tumors in a broad range of anatomical locations, including the head and neck region (oral mucosa, paranasal sinuses, tongue, salivary glands), thoracic cavity, stomach, kidney, ovary, skin, and limb soft tissues (Figure 5A), covering all reported anatomical locations of human NC. Most of these tumors were diagnosed as SCC with differing degrees of differentiation (Figure 5B and C and Supplementary Figure 5B and 5C), confirming a consistent phenotype. Notably, although the NLC-Cre driven mNCs consistently express NUTM1, cMYC and KRT14, the expression of P63 and SOX2 are variable (Figure 5B and C). The inter-tumor heterogeneity of these progenitor transcription factors is consistent with pathological observations in human patients and may be responsible for the remarkable plasticity and resistance to therapies of NC. Critically, atypical histological phenotypes were reported in isolated human NC cases, including glandular and chondrogenic metaplasia^53^. Consistently, regions of chondrogenic and glandular transitions were observed in some mNCs (Figure 5D). The new GEMM provided a critical tool for investigating the reprogramming capability of *Brd4::Nutm1* to drive NC across tissues with remarkable heterogeneity.

**Figure 5:**
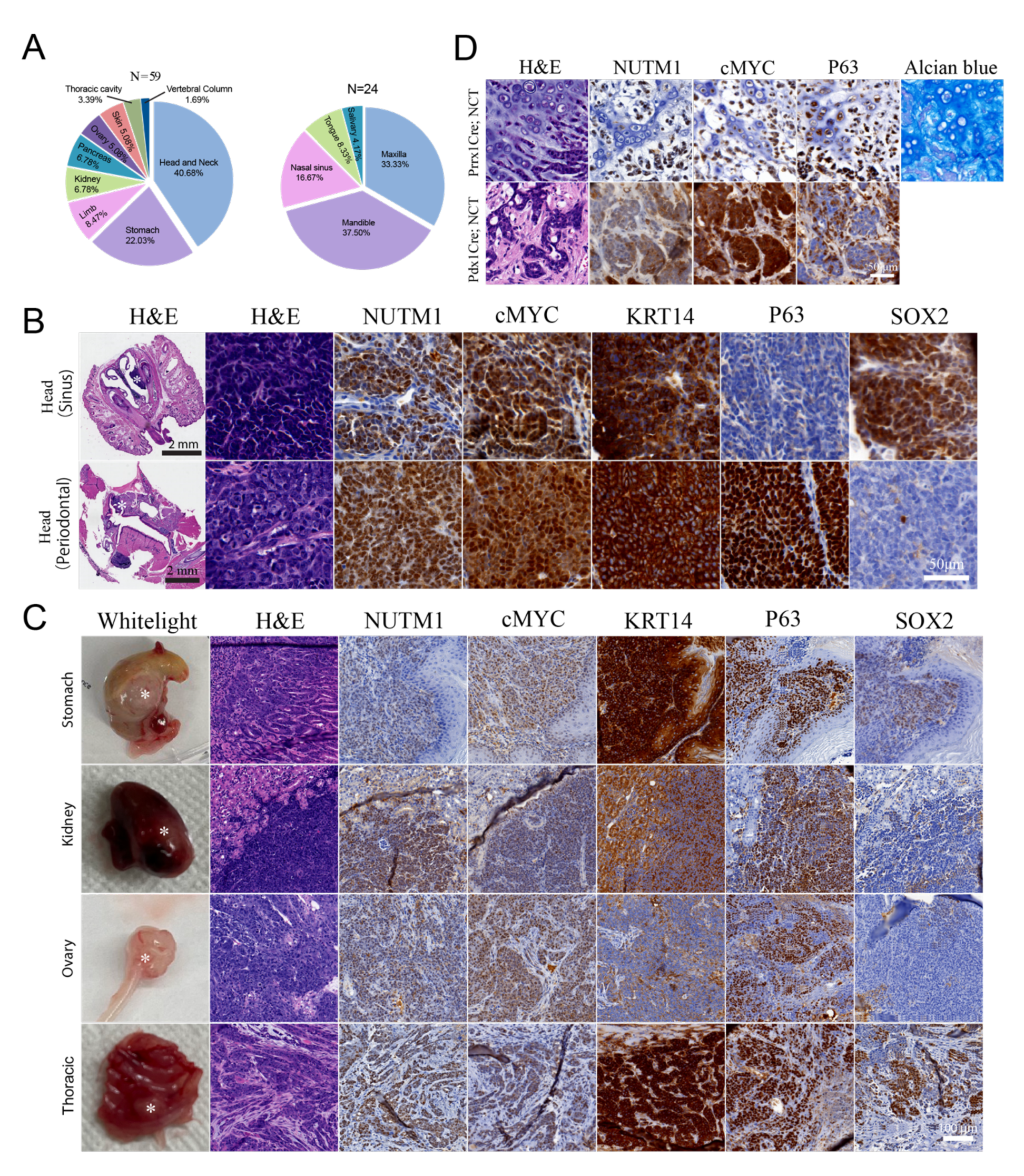
*Brd4::Nutm1* initiates NC across tissues. A. Tissue distribution of the NLS-Cre driven mNCs; left: distribution throughout body. Right. Distribution of the head and neck tumors. B. Representative whole-head H&E, high magnification H&E, and IHC images of nasal and oral NLS-Cre driven mNCs. C. Representative macroscopic, H&E, and IHC images of NLS-Cre driven mNCs outside the head and neck region. D. Representative H&E, IHC and Alcian Blue cartilage staining images of chondrogenic and ductal metaplasia in mNCs.

### 3.5. Mouse NCs have high metastasis potential

Most human NC patients have metastatic disease at diagnosis, contributing significantly to its dismal prognosis^2,54,55^. Of the more than 100 oral NC mice we analyzed, there were four cases of metastasis to proximal lymph nodes (Figure 6A), suggesting metastatic potentials of mouse NC. However, because the oral NCs lead to a quick death of mice, they may not allow the time for metastasis to establish and emerge. To further investigate, we focused on analyzing the Pdx1Cre driven pancreatic NC, which survives longer. Pancreatic NC mice consistently develop broad dissemination and metastasis at ten weeks of age. Large numbers of dissemination nodules (carcinomatosis) develop across the mesentery membrane (Figure 6B and Supplementary Figure 6A). Distant Micro-metastasis was detected in the lungs and liver (Figure 6B). The metastatic tumors express NC markers, including NUTM1, cMYC, P63, KRT14, and contain populations of SOX2-positive cells (Figure 6C).

**Figure 6:**
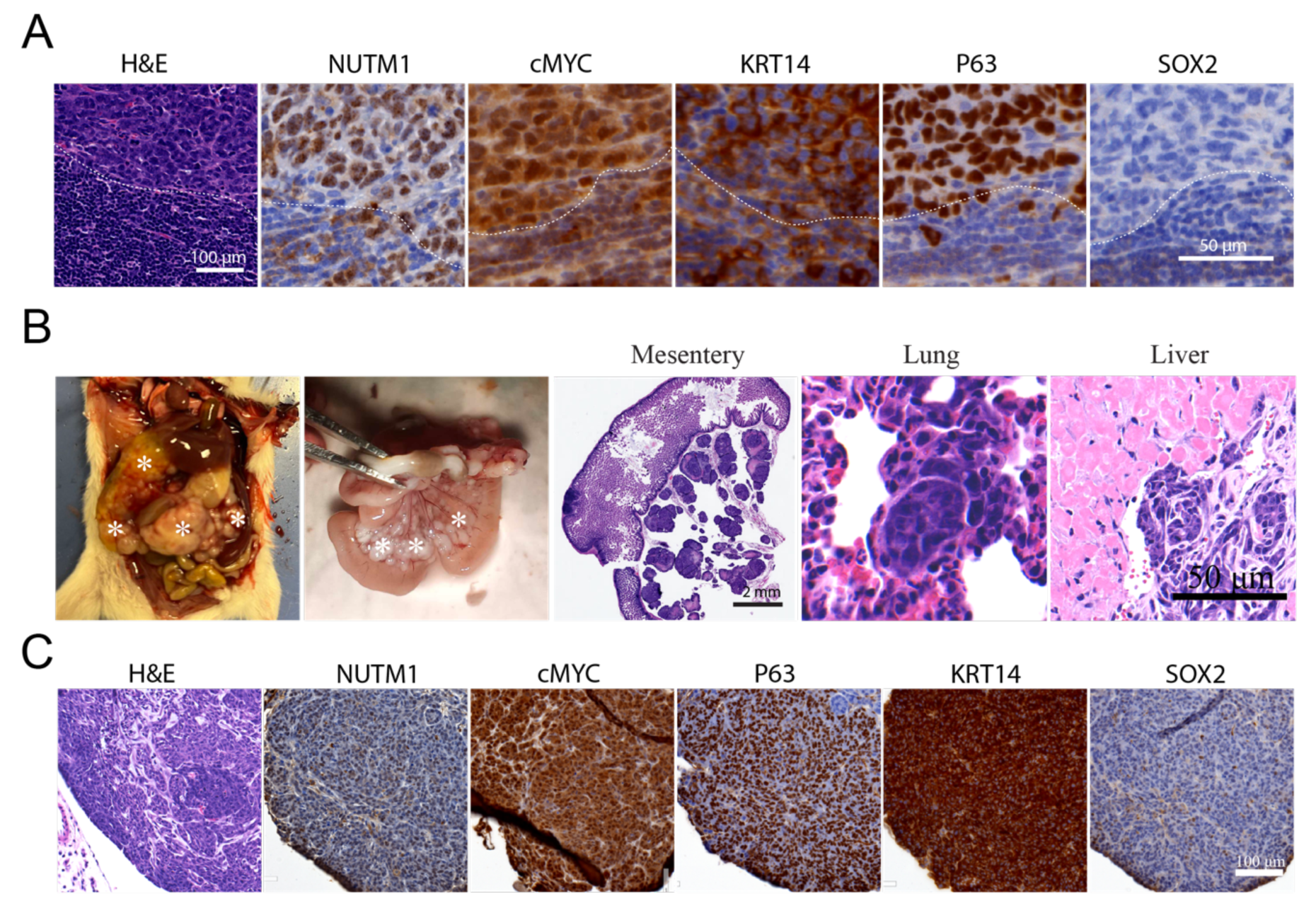
mNCs are highly metastatic. A. Representative H&E and IHC images of a lymph node metastasis from oral mNC. B. Representative macroscopic and H&E images of peritoneal/mesentry, lung and liver metastasis of pancreatic mNCs. C. Representative H&E and IHC images of peritoneal metastasis of pancreatic mNCs.

It is unclear why NCs are extremely metastatic. Desmoplasia, characterized by the recruitment of cancer-associated fibroblasts and pervasive deposition of extracellular matrix proteins, is strongly associated with metastatic potential in common cancers such as pancreatic ductal adenocarcinoma (PDAC)^56–58^. Desmoplastic histological features have been reported in a number of human NC case reports and are routinely observed in clinical diagnostic practice^59–61^, yet, have never been characterized at the molecular level. Through RNA sequencing analysis of mouse NC, we detected large increases in the expression of fibroblast genes such as *fibroblast activation protein (FAP)*, *smooth muscle alpha actin (Acta2)* and SM22alpha (*Tagln*) (Figure 7A). Mouse NC tissue also express high levels of ECM genes including *fibronectin (Fn1)*, *collagens* (17 types) and collagen cross-linking enzymes such as *lysyl oxidase (Lox)*, *Loxl2, Loxl3* and *peroxidasin (Pxdn)* (Figure 7B). Confirming these results, both mouse oral and pancreatic NC contain large numbers of activated fibroblast cells and abundant collagen-based ECM (Figure 7C). In addition to the unique desmoplastic stromal response, the epithelial-mesenchymal transition (EMT) process has been closely linked to metastasis^62^. A gene set enrichment analysis (GSEA) revealed significant activation of the Epithelial-mesenchymal transition program in mouse NCs (adjusted *p*-value = 2.16E-07 or 2.16x10^-7^), with strong upregulation of mesenchymal markers including Snai1^63^, Zeb1^64^ and Vimentin^65,66^ (Figure 7D). Consistently, at an advanced stage, regions of both mouse oral and pancreatic NC lost the membrane-localized epithelial marker E-Cadherin and gained the expression of Zeb1 and Vimentin to differing degrees (Figure 7E), demonstrating EMT, which might be associated with the metastatic potential of NC. It is clear now that EMT is a reversible process marking a continuous spectrum of phenotypes^67^. However, it is under debate whether full EMT is required for metastasis and whether a reverse process called Mesenchymal Epithelial Transition (MET) is required for the establishment of metastasis^62^. We analyzed EMT phenotypes of the metastatic lesions in mouse NC. As shown in Figure 7F, in some cases such as the lymph node metastasis of oral NC, the majority of cancer cells within the metastasis tumor expressed E-Cadherin but also vimentin and Zeb1, showing a mixed Epithelial-mesenchymal phenotype. On the other hand, the mesenteric nodules developed from pancreatic NCs have lost membrane localized E-cadherin but did not significantly upregulate Vimentin and ZEB1. Thus, there is likely a complex, non-binary interplay between desmoplasia, EMT, and metastasis of NC (more examples in supplementary Figure 6). The new GEMM provides a powerful tool to dissect these interplays.

**Figure 7:**
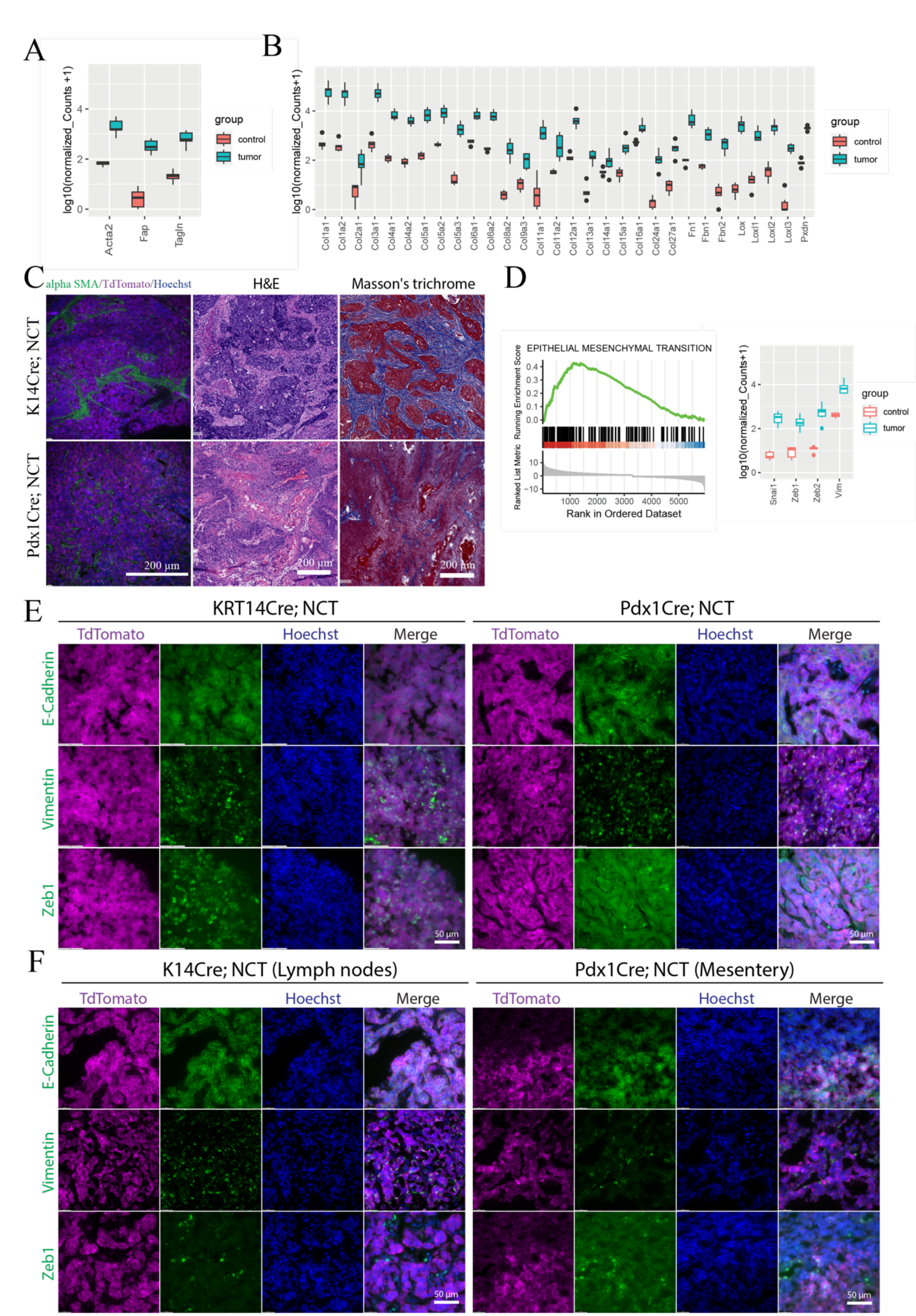
mNCs present desmoplastic and EMT features that could contribute to its metastatic potential. A. Upregulation of markers of activated fibroblasts in mNCs. B. Upregulation of ECM and their modifier genes in mNCs. C. Representative images of desmoplastic features of oral (upper) and pancreatic (bottom) mNCs. D. GSEA enrichment of EMT genes (left) and upregulation of key EMT markers in oral mNCs. E. Expression of the epithelial marker E-Cadherin and the Mesenchymal marker Vimentin and Zeb1 in advanced oral and pancreatic mNCs, showing regions of EMT. TdTomato staining demarcates NC cells. F. Expression of the epithelial marker E-Cadherin and the Mesenchymal marker Vimentin and Zeb1 in metastatic mNCs, presenting heterogenous EMT status. TdTomato staining demarcates NC cells.

## 3. Discussion

Nut carcinoma is an aggressive cancer with a dismal prognosis^2,10^. Because NC is extremely rare in the human population and is often diagnosed at a late disease stage, relevant *in vivo* models are vital for understanding its pathogenesis and for therapy development^20,25^. The new NC GEMM addressed key issues that are challenging to explore with currently available human cell line and xenograft models and created new opportunities for understanding NC.

First, a long-standing dilemma in NUT carcinoma research is that although the disease is strongly associated with chromosome translocations events that induce the expression of NUTM1 fusion genes such as *BRD4::NUTM1*, the ectopic expression of the fusion genes in non-NC cells are unequivocally cytotoxic^2,24^. This dilemma raises a critical question as to whether the *NUTM1* fusion genes are indeed responsible for NC initiation. Through analyzing hundreds of NC GEMM, our data demonstrated that when expressed at the endogenous level from an endogenous chromosome translocation, *BRD4::NUTM1* can efficiently induce aggressive cancers with high penetrance. The mouse tumors that BRD4::NUTM1 induced presented similar histopathological and molecular features of human NCs. mNCs also reproduced the intrinsic intra and inter-tumor heterogeneity. This comprehensively characterized GEMM firmly demonstrates that the *BRD4::NUTM1* fusion gene is sufficient to induce NC-like tumors *in vivo*.

Second, although human NC is rare, cases have been reported in a broad range of tissues. The main affected sites were reported to be head and neck (∼40% of reported cases) and thoracic (∼50% of reported cases)^3^. There are two possible explanations for the predilection for these sites: 1) The cell types in these tissues are more receptive to transformation by *BRD4::NUTM1*. 2) Fewer cases were diagnosed at other sites due to limited clinical awareness. The second explanation is not trivial, considering the history of NUT carcinoma. It was first thought to be a pediatric thymic cancer^1^. With the increased awareness and better availability of diagnostic methods such as the NUTM1 IHC and next-generation sequencing, NC is now considered a preferentially young adult cancer that can occur in any age group and many tissue sites^3^. Our data showed that when *Brd4::Nutm1* fusion was induced in tissue progenitors of all three germ layers using cell type-specific cre drivers, NC can be induced at 100% penetrance in the respective tissue. Furthermore, when *BRD4::NUTM1* fusion was induced stochastically using a constitutively active Cre driver, mNCs can arise in many tissues, such as the paranasal sinuses, ovary and kidney, previously reported in humans, and previously not reported tissues, such as the gastric tissues. Our data raised a critical possibility that the low case rate outside the head and neck/thoracic region might be a result of diagnosis bias, either due to limited clinical awareness or in the case of gastric cancer, the low general prevalence and research intensity in North America^68^. Broader and global surveys of NUT Carcinoma cases in hard-to-diagnose cancers with monolithic, poorly differentiated morphology in all tissues may establish a more complete picture of NC in humans. At the molecular level, the broad cell type receptivity and the universal SCC phenotype implied that BRD4::NUTM1 coerces a shared molecular program in cell types across all germ layers to initiate NC. Yet, BRD4::NUTM1 has a strong reprogramming ability to drive an aggressive poorly differentiated SCC phenotype of NC regardless of lineage context. The new GEMM provided a unique model to further investigate the key molecular mechanisms underlying NC oncogenesis.

Fourth, human NCs are highly metastatic, yet, currently, there is no established model to study this key aggressive phenotype. The GEMM uncovered an extremely strong metastatic potential of NC. GEMMs are not always optimal for studying metastasis. In many GEMMs of common cancers, mice reach their endpoint due to the growth of primary tumors before significant metastasis^69^. In GEMMs that consistently develop metastasis, such as the mutant K-Ras driven pancreatic ductal carcinoma (PDAC) model, metastasis takes a long time to develop (15-40 weeks) and is challenging to study^49,70^. The pancreatic NC GEMM develops widespread metastasis in a short time window (5-10 weeks), providing ample materials to investigate the mechanisms of NC metastasis in vivo. Critically, the primary mouse NCs do present phenotypes, including desmoplasia and EMT that are associated with metastatic potentials. Further investigation using the new GEMM will lead to a fruitful understanding of the metastasis mechanism and therapeutic strategies of NC.

As immune therapy has transformed the therapeutic landscape of aggressive cancers such as melanoma and lymphoma, it is critical to investigate the immune system’s interaction with targeted or immune therapies for NC. The GEMM reported here is the first immune-competent experimental model for NC and can provide a valuable platform for studying the immune therapies for NC.

After our preprint was online, another NC mouse model was submitted and published^71^. This model uses the *cre*-dependent one-way genetic switch (FLEx) conditional inversion system and a Sox2-CreERT2 tamoxifen driver to induce the *Brd4::Nutm1* fusion gene from the mouse Brd4 allele. In parallel, these two GEMM each have unique advantages and disadvantages in modeling NC. First, the FlEx model is driven by Tamoxifen inducible Cre, thus have better temporal control. However, the high recombination efficiency will mean that the fusion gene will be induced in a large number of cells simultaneously, mimicking the sequencing mutagenesis and field carcinogenesis in common cancers^72^, different from the clonal stochastic reality of fusion gene driven cancer. Our translocation model recapitulates such a reality.

Second, the FlEx model induces mNCs only in the esophagus, providing a clear and easily accessible target tissue for study. However, there has been no report of primary NC in the esophagus. Whether this inconsistency is due to species difference, miss- assigning of human primary NC, or an artifact due to pervasive induction of *Brd4::Nutm1* in all esophagus basal cells by the Sox2-CreERT, remains to be determined. In contrast, our translocation model closely recapitulated the human tissue spectrum, including the rare cases outside the head-and-neck and thoracic regions, providing a richer resource understanding of the NC oncogenesis from different tissues. Furthermore, the broad tissue distribution of the translocation model will allow intersectional analysis to find the core mechanisms that drive NC and model more tissue contexts, which will be critical for future drug delivery and immune therapy research.

In summary, we have reported the first GEMM for NC that recapitulates the endogenous chromosome translocation. The new GEMM can enhance our understanding of this disease and open up many opportunities for developing targeted therapies. The GEMM will serve as a valuable preclinical tool for the global community to study the mechanism of NC and develop new treatments.

## 4. Materials and Methods

### 4.1 Ethical statement

All animal work was carried out under PROTO202000143 and PROTO 202300127, approved by the Michigan State University (MSU) Campus Animal Resources (CAR) and Institutional Animal Care and Use Committee (IACUC) in AAALAC credited facilities.

### 4.2 Mouse line generation

All mouse generation work follows our published 2-cell (2C)- Homologous Recombination (HR)- CRISPR (2C-HR-CRISPR) method and protocol and is described briefly as follows^34,73,74^. First, 2-cell stage embryos of the CD1 strain were microinjected with Cas9-mSA mRNA (75ng/ul), sgRNA targeting around the stop codon of the Nutm1 gene (GGUGGCCCUCUGCUUCCUAC, chemically modified sgRNA from Synthego Inc. 50ng/ul) and a biotinylated PCR donor consisted of homology arms (5’arm 1000bp and 3’arm 600bp) spanning the coding sequence of T2A-Luc2TdTomato cassette (5ng/ul).

The Luc2TdTomato cassette was cloned from the pCDNA3.1(+)/Luc2-TdT plasmid (Addgene 32904, kind gift from Christopher Contag)^35^. Injected embryos were implanted into pseudopregnant females the same day to generate founder pups. Positive founders were screened by long-range PCR spanning homology arms using the following primer sets. 5’ arm gtF: GAGACTGTCATAGACAGCATCCAAGAT and 5’arm gtR: tcatggctttgtgcagctgc; 3’arm gtF: ggactacaccatcgtggaacagta and 3’arm gtR: CATATTTAACAGTCCCACGGAGAG. A positive founder mouse was out-crossed to CD1 mice to generate N1 mice. The N1 mice were genotyped by PCR. Additionally, genomic regions spanning the targeting cassette and 3’ and 5’ homology arms were Sanger-sequenced to validate correct targeting. Insertion copy number was evaluated by digital droplet PCR ddqPCR. Heterozygous N1 mice have only one insertion copy, demonstrating single-copy insertion. An N1 that passed all quality controls was further outcrossed with CD1 for two generations to remove any potential off-target mutations introduced by CRISPR-Cas9 editing to generate N3. Heterozygous N3s were bred together to establish a homozygous colony of the Nutm1-T2A-Luc2TdTomato mousseline.

Then two additional mouse lines were generated in parallel. First, a loxp site was inserted into the intron 10 of the Brd4 gene (Brd4loxp) by electroporating CD1 zygotes with Cas9 proteins, a chemically modified sgRNA (GAAUAGAGUGCUGCUACACU, Synthego Inc.), and a single strand oligo DNA donor (ssODN), with 30 base homology arms on each side spanning the loxp site (TCTCCCTGCAGAGTGGCAAGATGCCCAGTGataacttcgtatagcatacattatacgaagttatTAGCAGCACTCTATTCTCTCCATCTGGGTA). Electroporated zygotes were implanted into pseudopregnant females the same day to generate founder pups. Positive founder pups were screened by PCR using primers: BRD4 loxp gt F: TGCCTGGGCTCTTCTGTCCAC, and BRD4 loxp gt R: AACCCAAGGTTGATGGGACCC. A positive founder mouse was out-crossed to CD1 mice to generate N1 mice. The N1 mice were genotyped by PCR. Additionally, genomic regions spanning the targeting cassette and 3’ and 5’ homology arms were Sanger-sequenced to validate correct targeting. An N1 that passed all quality controls was further outcrossed with CD1 for two generations to remove any potential off-target mutations introduced by CRISPR-Cas9 editing to generate N3. Second, a loxp site was inserted into the intron 1 of the Nutm1-T2A-Luc2TdTomato gene (Nutm1-loxp) by microinjecting Nutm1-T2A-Luc2TdTomato homozygous 2-cell embryos. (75ng/ul), sgRNA targeting around the intron (ACCGGAUCCCAACCGCUCUA, chemically modified sgRNA from Synthego Inc. 50ng/ul) and a biotinylated PCR donor consisting of homology arms (5’arm 500bp and 3’arm 500bp) spanning a loxp site and artificial genotyping primer sequences (5ng/ul). Injected embryos were implanted into pseudopregnant females the same day to generate founder pups. Positive founders were screened by long-range PCR spanning homology arms using the following primer sets. 5’ arm gtF: GCATGAAATCAGACCAAGTGGGC and 5’arm gtR: AAGCTGACCCTGAAGTTCATCTG; 3’arm gtF: ggactacaccatcgtggaacagta and 3’arm gtR: CATATTTAACAGTCCCACGGAGAG. A positive founder mouse was out-crossed to CD1 mice to generate N1 mice. The N1 mice were genotyped by PCR. Additionally, genomic regions spanning the targeting cassette and 3’ and 5’ homology arms were Sanger-sequenced to validate correct targeting. An N1 that passed all quality controls was further outcrossed with CD1 for two generations to remove any potential off-target mutations introduced by CRISPR-Cas9 editing to generate N3. Starting from N3s, the Brd4loxp and Nutm1loxp strains were bred to double homozygous to generate the NC translocator (NCT) mouse line.

For inducing the BRD4::NUTM1 fusion, NCT+/+ mice were crossed with heterozygous KRT14Cre mice (Jackson Lab 018964) for oral cancer, Pdx1Cre mice (Jackson Lab 014647) for pancreatic NC, Prrx1Cre mice(Jackson Lab 005584) for limb soft tissue NC and NLS-Cre (MGI:3586452) for universal screen. The Cre mouse lines from Jackson Lab was on the B6 background and had black fur. To facilitate sensitive bioluminescent imaging, the strain was outcrossed for two generations with CD1 mice to generate albino mice. Progenies from the crosses were genotyped; the cre-positive ones were used as experimental groups, while the cre-negative ones were used as controls.

#### Bioluminescence imaging

NC mice were subjected to regular bioluminescence imaging to monitor tumor signals. During the imaging procedure, mice received an intraperitoneal injection of 150 mg/kg D-Luciferin Potassium (Goldbio). After 20 minutes, images of both dorsal and ventral positions were captured using IVIS Spectrum In Vivo Imaging System (Perkin Elmer) under anesthesia. To quantify the tumor signal intensity, regions of interest (ROIs) were delineated around the head area. The total flux (photons/sec) within the ROIs was measured using the Living Image® Software. The average value of the total flux from both dorsal and ventral images was used for generating tumor growth curves.

### 4.3. Necropsy and tissue collection

Routine necropsies were performed by ventral midline approach. Tumors, draining lymph nodes, heads, lungs, and liver were harvested and fixed in 10% neutral buffered formalin for 24 h or flash-frozen in liquid nitrogen for further analyses. Tissue samples were either fixed in 4% paraformaldehyde at 4° C for 24 h or 10% neutral buffered formalin at room temperature for 24 h. Heads were decalcified in 10% EDTA for 12-14 days at 4° C by replacing the decalcification solution every two days. Tissues were either routinely processed, and embedded in paraffin wax or used for cryo-sections.

### 4.4. Histopathology

Serial, 5-µm thick sections were cut and stained with hematoxylin-eosin (H&E). Neoplasms were characterized based on anatomic location, necrosis, inflammation (polymorphonuclear and mononuclear), and histologic features (focal dysplasia, undifferentiated carcinoma, poorly differentiated carcinoma with regional squamous differentiation, and well-differentiated squamous cell carcinoma as described^75^. Histopathologic examination was performed by a board-certified veterinary pathologist (MFT).

### 4.5. Immunohistochemistry

Antigen retrieval was done using sodium citrate pH 6 for 20 min at 97 °C. Immunohistochemistry was performed using antibodies targeting NUTM1 (1:100, Huabio HA721690), TdTomato (1:500, Origene, ABOO40-500), cMYC (1:500, Proteintech, 10828-1-AP), SOX2 (1:1000, R&D Systems, AF2018), TP63 (1:1000, Sigma Aldrich, AMAB91224), KRT14 (1:500, Novus Biologicals, NBP2-67585), and involucrin (1:1000, BioLegend, 924401) using Agilent Dako Autostainer Link 48 and EnVision FLEX kit (Dako). Immunoreactivity for each antibody was assessed by light microscopy.

### 4.6. Cryosection

Tissues for cryosections were dehydrated using 30% sucrose at 4° C overnight. Samples were embedded in OCT and snap-frozen on a steel bench block immersed in liquid nitrogen. Samples were sectioned at 4-µm thickness using a cryostat (Leica).

### 4.7. Immunofluorescence

Slides were dried at room temperature for 30 minutes and permeabilized using 0.1% Tween 20 for 30 minutes. Samples were blocked using PBST containing 5% normal donkey serum for 1 hour at room temperature. The primary antibodies (same as described in the IHC section) diluted in blocking solution were applied on tissue sections and incubated overnight at 4 °C. Slides were rinsed three times for 5 minutes each with PBS. Secondary antibodies diluted in blocking solution were applied on tissue sections and incubated in a dark humid chamber for 1 hour at room temperature. Slides were rinsed three times for 5 minutes each with PBS. Nuclei were stained with Hoechst 33258 and then mounted with Vectashield® mounting medium. Slides were imaged with a Leica Thunder microscope imaging system.

### 4.8. Duo color FISH

The Oligopaint FISH probe libraries were constructed as described previously (Xie et al., Nature Methods, 2020). A ssDNA oligo pool was ordered and synthesized as custom arrays from GenScript. Each oligo consists of a 30-37 nucleotide (nt) homology to the 30kb region, flanking the mBrd4-mNut1m fusion point from the mm10 genome assembly. Each Oligo was run through the OligoArray2.0 with the parameters -n 22 -D 1000 -l 32 -L 32 -g 52 -t 75 -T 85 -s 60 -x 60 -p 35 -P 80-m "GGGGG;CCCCC; TTTTT;AAAAA" according to algorithms developed from the Wu laboratory (https://oligopaints.hms.harvard.edu/). The mBrd4 and mNut1m oligo subpool consists of a unique set of primer pairs for PCR amplification, a 20 nt T7 promoter sequence for in vitro transcription, and a 20 nt region for reverse transcription. The mBrd4 oligo pool was amplified by GCGGGACGTAAGGGCAACCG and GCGTTGCGGTGCGATCTTCT.

The mNut1m was amplified by TGATAACCCACGCACGGCTG and GACCCGGGCCACTAACACGA. Following PCR amplification, each Oligopaint probes were generated by in vitro transcription and reverse transcription, in which ssDNA oligos conjugated with ATTO565 and ATTO647 fluorophores were introduced during the reverse transcription step. The Oligopaint covered genomic regions (mm10) used in this study are listed below: mBrd4, chr17:32,210,998-32,244,827. mNut1m, chr2:112,227,883-112,257,004. The oligo sequences for conjugating fluorophores during reverse transcription are: /5ATTO565N/AATTCGGCAGACCCGAATGC (mNut1m) and /5ATTO647N/CCTATCCCCTGTGTGCCTTG (mBrd4).

DNA FISH is performed on cryosections prepared as described above. Tissue slides were washed 2X with 1XPBS and permeabilized in 0.5% TritonX100 in 1XPBS for 60min in a Coplin Jar. After 2X wash in 1XPBS, slides were treated with 0.1M HCl for 5min, followed by 3X washes with 2XSSC and 30 min incubation in 2X SSC + 0.1% Tween20 (2XSSCT) + 50% (v/v) formamide (EMD Millipore, cat#S4117). For each sample, we prepare a 25ul hybridization mixture containing 2XSSCT+ 50% formamide +10% Dextran sulfate (EMD Millipore, cat#S4030) supplemented with 0.5µl 10mg/mL RNaseA (Thermo Fisher Scientific, cat# 12091021) +0.5µl 10mg/mL salmon sperm DNA (Thermo Fisher Scientific, cat# 15632011) and 20pmol probes with distinct fluorophores. The probe mixture was thoroughly mixed by vortexing and briefly microcentrifuged. The hybridization mix was transferred directly onto the slide covered with a clean coverslip. The coverslip was sealed onto the slide by adding a layer of rubber cement (Fixo gum, Marabu) around the edges. Each slide was denatured at 80°C for 5 min, then transferred to a humidified hybridization chamber and incubated at 42°C for 16 hours in a heated incubator. After hybridization, samples were washed 2X for 15 minutes in prewarmed 2XSSCT at 60 °C and then were further incubated at 2XSSCT for 10min at RT, at 0.2XSSC for 10min at RT, at 1XPBS for 2X5min with DNA counterstaining with DAPI. Then coverslips were mounted on slides with Prolong Diamond Antifade Mountant (Thermo Fisher Scientific Cat#P36961) for imaging acquisition.

DNA FISH images were acquired using a structured illumination microscopy on a Zeiss AxioObserver microscope (Zeiss Elyra7). Images were taken with a Plan-Apochromat 63x/1.40 oil DIC objective in a lens immersion medium having a refractive index of 1.515 and two pco.edge 4.2 CL HS sCMOS cameras. We used 405nm (Excitation wavelength) and 460nm (Emission wavelength) for the DAPI channel, 561nm (Excitation wavelength) and 579nm (Emission wavelength) for the ATTO565 channel, and 633nm (Excitation wavelength) and 654nm (Emission wavelength) for the ATTO647 channel. We acquired 13 phase images for each focal plane. Image post-processing was performed with the ZEN 3.0 SR (black) software.

### 4.9. RNA sequencing

Total RNA was prepared from tumors and normal tissues using the RNeasy Mini Kit (Qiagen Inc.). The quality control, cDNA library preparation and bulk RNA sequencing were performed by Novogene Inc following standard protocol using the Novaseq 6000 sequencer. Each sample were sequenced for 20M reads.

FastQC v0.11.7^76^ was used to evaluate sequence quality, GC content, overrepresented sequences and adapter contamination of the raw paired-end FASTQ reads for each sample. Trimmomatic v0.39^77^ was used to trim raw reads using following settings: PE ILLUMINACLIP: TruSeq3-PE.fa:2:30:10:8:true HEADCROP:15 CROP:110. All samples passed quality control based on the results of FastQC. Quality trimmed reads were pseudoaligned to GENCODE mouse transcript reference (GRCm39, release M32) and transcription levels were quantified using kallisto^78^ (version: 0.46.1, parameters: -b 100 - l 110). Gene expression levels were generated using tximport v1.28.0^79^ and filtered for lowly expressed genes (0.5 count per million in at least six samples). Differential expression analysis were carried out using the Bioconductor v3.17.1^80^ package DESeq2 v1.40.2^81^. rlog-normalized counts were obtained from the differential expression analysis and used for principal component analysis, sample correlation calculation and k-means clustering^81^. The Wald test was used by DESeq2 to identify genes that are differentially expressed. Benjamini-Hochberg was used to adjust the p values for multiple testing^82^.

The log2 fold change (LFC) was shrunken by using the “apeglm” method to provide more accurate estimates^83^. Differentially expressed genes (DEGs) were defined based on LFC > 1 or < -1 and adjusted *p*-value < 0.05. ClusterProfiler v4.8.2^84^ was used to perform functional enrichment analysis by using different databases: Gene Ontology^85,86^, KEGG PATHWAY^87^, Reactome^88^ and the Hallmark gene sets of Molecular Signatures Database (MSigDb)^89^. Lists of downregulated DEGs and upregulated DEGs were separately examined for Over Representation Analysis (ORA)^90^. The shrunken LFCs were used to rank all DEGs decreasingly to generate a gene list as an input for Gene Set Enrichment Analysis (GSEA)^91^. Redundant GO terms were removed by GOSemSim with a cut-off of 0.7^92^. Pathways or gene sets with adjusted p value < 0.05 were considered significantly enriched. RNA-seq data will be available through GEO after publication.

### 4.10. Whole genome sequencing

Genomic DNAs was extracted from normal and tumor samples using The GeneJet DNA kit (Thermo Fisher), according to the manufacturer’s instructions. The quality control, cDNA library preparation and RNA sequencing were performed by Novogene Inc following standard protocol. A total of 1.0 ug DNA per sample was used for library preparations. Each sample were sequenced for 10-16X coverage. WGS data will be available through GEO after publication.

Raw reads for each sample were trimmed using Trimmomatic v0.39^77^ (PE ILLUMINACLIP:2:30:10:8:true HEADCROP:5 CROP:120). Then the trimmed reads were aligned to the GENCODE mouse primary genome assembly (GRCm39, release M32) using the BWA-MEM algorithm in the Burrows-Wheeler Aligner (BWA) v0.7.17^93^ with following options: -M to flag shorter split hits as secondary; -R to add read group information. The SortSam tool of Picard v2.25.0^94^ was used to sort the output files from BWA by queryname and export output as binary Sam (BAM) files. Subsequently, duplicate reads were marked using markDuplicates tool in Picard and the resulting files were sorted using SortSam by coordinates. The AlignmentSummaryMetrics and CollectWgsMetrics tools of Picard were used to collect alignment and genome coverage statistics. The Mutect2 tool of GATK v4.1.4.1^95^ used to identify and call variants in each sample individually. Variants were firstly filtered using FilterMutectCalls of GATK to remove probable technical or germline artifacts. Additional filters can be used to decrease the false-positive rate via SnpSift v4.1^96^: mutant allele frequency (≥5%), coverage at particular positions in tumor and normal samples (≥5×) and supporting reads for the mutation in the tumor samples (at least two). Filtered variants were further compared to exclude known polymorphisms by using bcftools v1.9.64^97^. snpEff v4.1^96^ was used to anotate the effects of variants. Large structural variations were identified using Delly v0.7.8^98^.

### 4.11. Cell-type deconvolution analysis

The signature matrixes were separately generated from two published single cell RNA-seq datasets for implementation in the CIBERSORTx deconvolution algorithm^99^. Briefly, the preprocessed count matrix of mouse tongue from 10x genomics sequencing was extracted from the Tabula Muris. Quality control, normalization, and clustering were performed as described in the original study^46^. The transcripts per kilobase million (TPM) gene expression profiles of 7538 labeled cells, pertaining to 7 distinct cell types (proliferating basal cells, basal cells, suprabasal differentiating keratinocytes, suprabasal differentiated keratinocytes, differentiated keratinocytes, filiform differentiated keratinocytes and Langerhans cells), were used to build a signature matrix file (q value = 0.1). The number of barcode genes per cell type was held between 300 and 500, resulting in a matrix consisting of 2256 genes. With the same strategy, the preprocessed, mouse oral epithelial basal layer cells-derived single-cell RNA-seq dataset from Jones KB, et al.^47^ was also used to generate a signature matrix of with 8 cell subtypes and 1970 genes included. The two signature matrixes were then applied to deconvolute the bulk RNA-seq TPM matrix of mNC tumor and normal control to predict the fraction of cell subtypes, with the absolute mode, S-mode batch correction, and 100 permutations.

## Karyotyping

Karyotyping analysis was performed on two early passage cell lines derived from mouse oral NC by Karyologic Inc. following standard protocol.

## Software and statistics

The tumor growth and survival data were analyzed and visualized using Prism 9 software (GraphPad, USA). Animal sacrifice was considered the endpoint for both analyses. The median survival times were calculated utilizing the Kaplan-Meier analysis, and significance values were determined by conducting the log-rank test. A p-value of less than 0.05 defined statistical significance.

Author contribution:

D.Z: formal analysis, investigation, methodology, validation, visualization, writing review, and editing;

A.E: formal analysis, investigation, methodology, validation, visualization, writing review and editing;

C.L: formal analysis, investigation, methodology, validation, visualization, writing review and editing;

M.B: methodology and visualization for FISH

L.X: methodology and visualization for FISH

D.B: Pathology analysis, human pathology insights, writing, review and editing; Y.T: Human clinical insights, writing, review and editing;

E.H: Human clinical insights, writing, review and editing; G.M: Bioinformatics analysis, writing, review and editing;

M.T: histopathological and IHC analysis, writing, review and editing;

B.G: conceptualization, design and generation of NC GEMM, formal analysis, investigation, methodology, validation, visualization, writing original draft, writing review and editing; providing funding support;

## Acknowledgment

We thank Dr. Andrew Sikora, Dr. Xiao Zhao and Dr. Jennifer Wang from the MD Anderson Cancer Center for constructive discussions. We thank MSU Transgenic and Genome Editing Facility (TGEF) for their help in generating the NC GEMM and the Histology laboratory at Veterinary Diagnostic Laboratory (Taylor Vaughn) for their help in histopathology and IHC analysis. We would also like to thank MSU Precision Health Program Tissue Analysis Core for assistance in slide scanning.

---

**Supplementary Figure 1.** The validation of the NC GEMM. A. Representative BLI images of NCT mice. B. A representative IF image of NUTM1 and TdTomato in the testis of NCT mice. C. A representative IF image of TdTomato in a whole head section of Krt14Cre; R26-LSL-TdTomato mice, depicting the tissue distribution of Cre activities. D. BLI images of 19 representative Krt14Cre-dirven NC mice. E. The tissue distribution of Krt14Cre-dirven mNCs. F. A representative sange sequencing track of the *Brd4::Nutm1* fusion junction. G. A representative karyotype of Krt14Cre-dirven mNC cells. Arrows indicates the t(2;17) translocation. H. Circos chart of chromosome translocation detected by whole genome sequencing of two mNCs. The thick blue line indicates t(2;17) forming *Brd4::Nutm1* is the only recurrent chromosome translocation. I. Two additional whole head NUTM1 IHC in mNC bearing mice.

**Supplementary Figure 2.** Molecular features if mNCs. A. A representative IHC image showing NUTM1 nuclear foci in mNCs. B. Representative IF images showing H3K27ace nuclear foci in mNCs. C. Representative whole head section IHC images of P63, KRT14, cMYC and SOX2 in two mNC bearing mice.

**Supplementary Figure 3.** RNA sequencing analysis of mNCs. A. RNA integrity number of samples demonstrated appropriate quality. B. Clustering analysis of RNA sequencing data. C. Principal Component Analysis (PCA) of RNA sequencing data D. CIBERSORT deconvolution of the bulk RNA sequencing data using a second set of single-cell RNA sequencing signatures of oral mucosa epithelium.

**Supplementary Figure 4.** BRD4::NUTM1 induced pancreatic and soft tissue mNCs. A. BLI images of 18 Pdx1Cre;NCT mice. B. Representative H&E and IHC images of pancreatic NC from Pdx1Cre;NCT mice. C. BLI images of 9 Prrx1Cre;NCT mice. D. B. Representative H&E and IHC images of soft tissue NC from Prrx1Cre;NCT mice.

**Supplementary Figure 5.** BRD4::NUTM1 induced NC from a broad range of tissues. A. BLI images of 25 NLS-Cre; NCT mice. B. Histopathological diagnosis of 62 mNCs derived from NLS-Cre; NCT mice. C. Additional representative H&E images of mNCs derived from NLS-Cre; NCT mice. D. Additional representative H&E images of metaplastic mNCs derived from NLS-Cre; NCT mice.

**Supplementary Figure 6.** Metastatic lesions from pancreatic mNC. A. Representative macroscopic images of metastatic mNCs. B. Representative IF of EMT markers of abdominal metastasis from pancreatic mNCs. C. Representative IF of EMT markers of diaphram metastasis from pancreatic mNCs. D. Representative IF of EMT markers of thymic metastasis from pancreatic mNCs.

