## Supplementary figures and images for "*Brd4::Nutm1* fusion gene initiates NUT carcinoma *in vivo*"

### Supplementary Figure 1

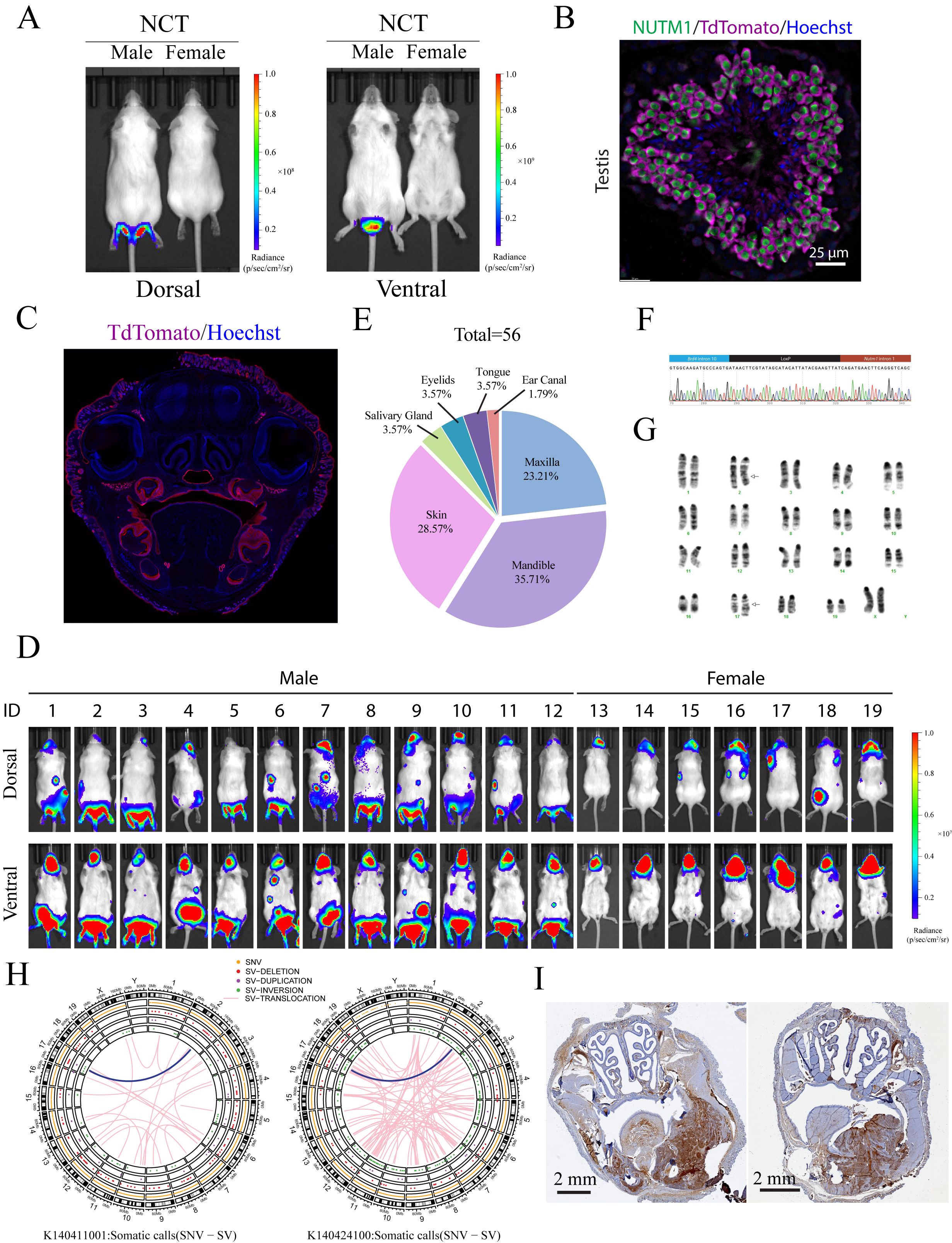

### Supplementary Figure 2

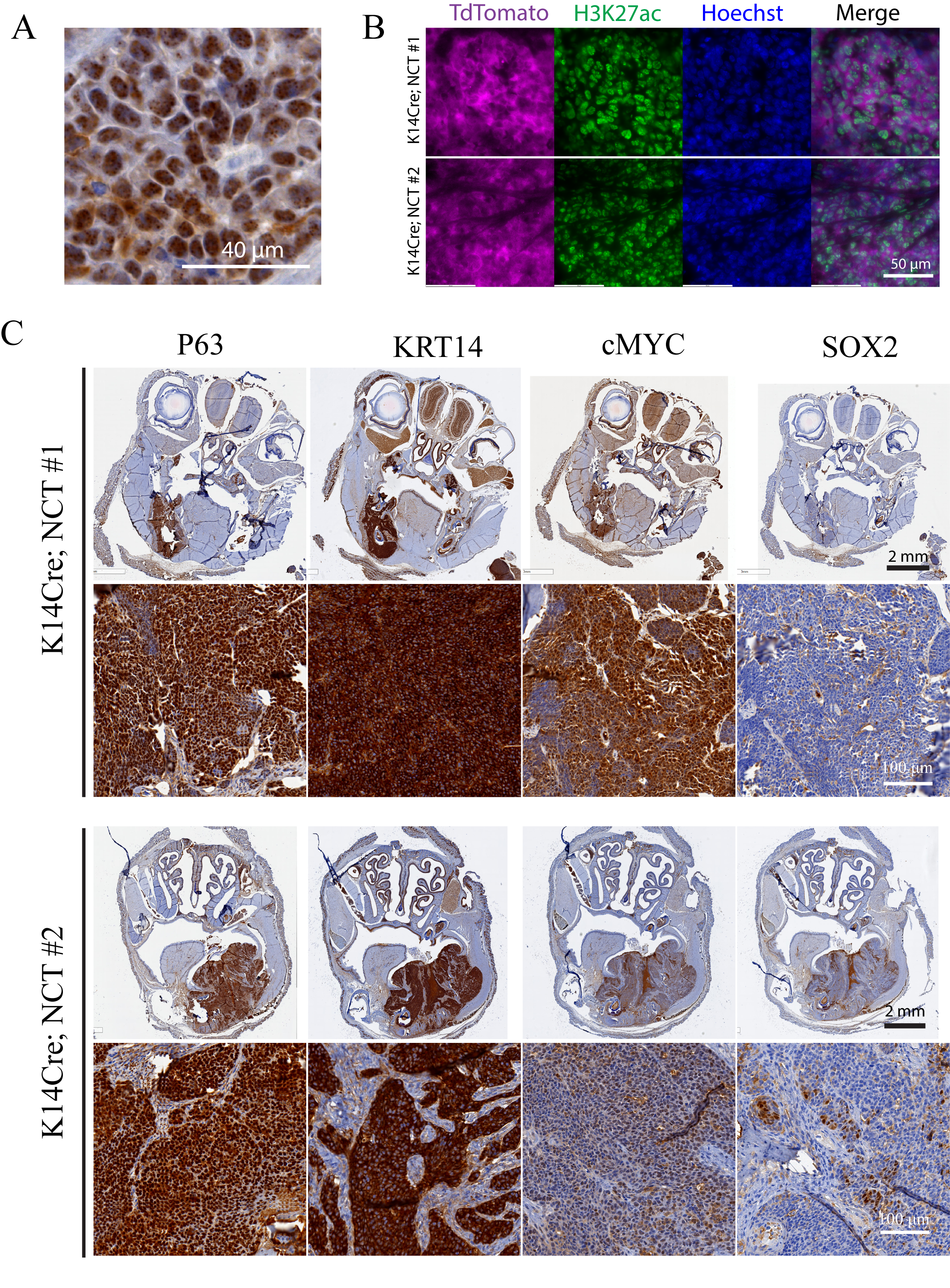

### Supplementary Figure 3

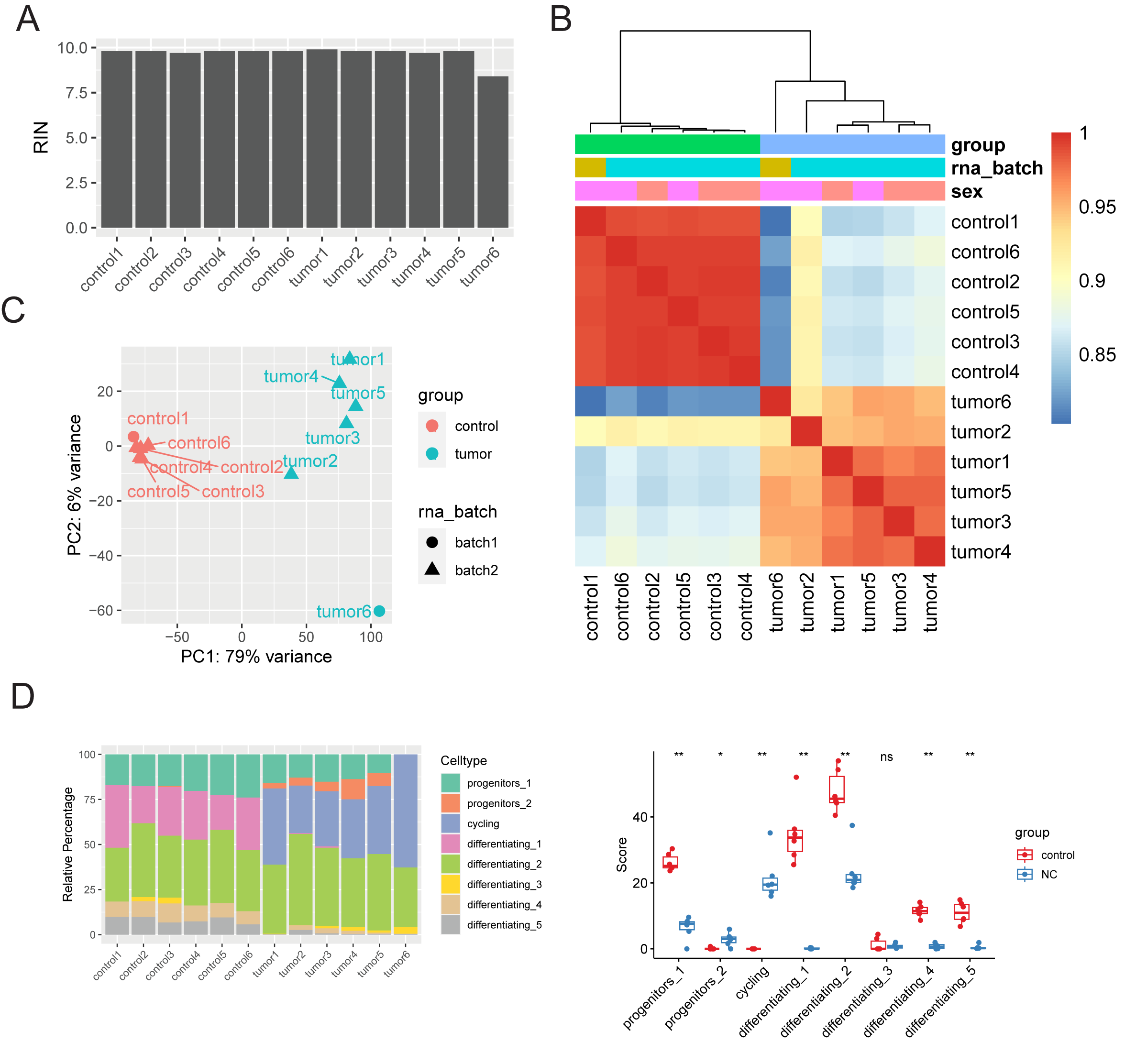

### Supplementary Figure 4

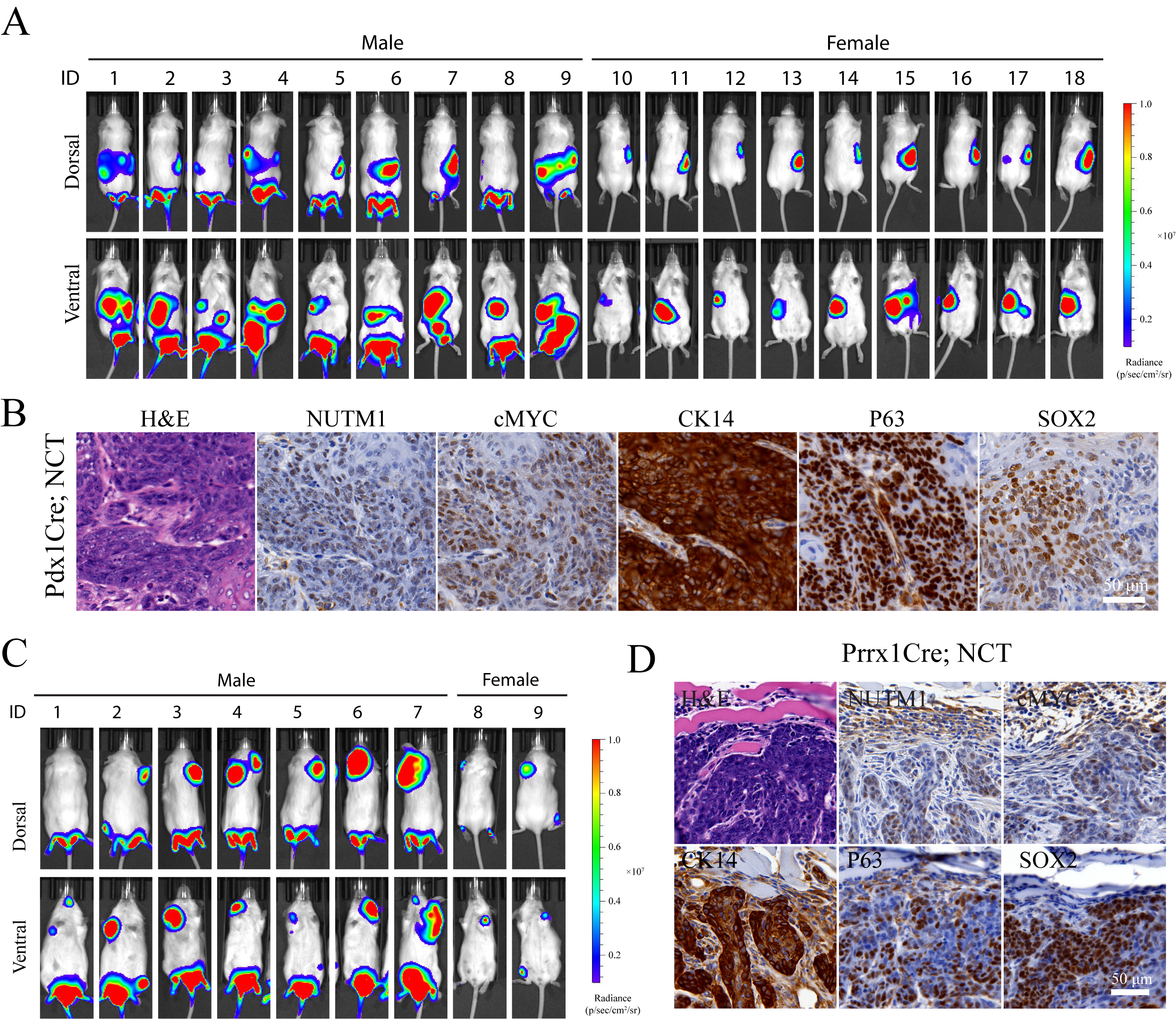

### Supplementary Figure 5

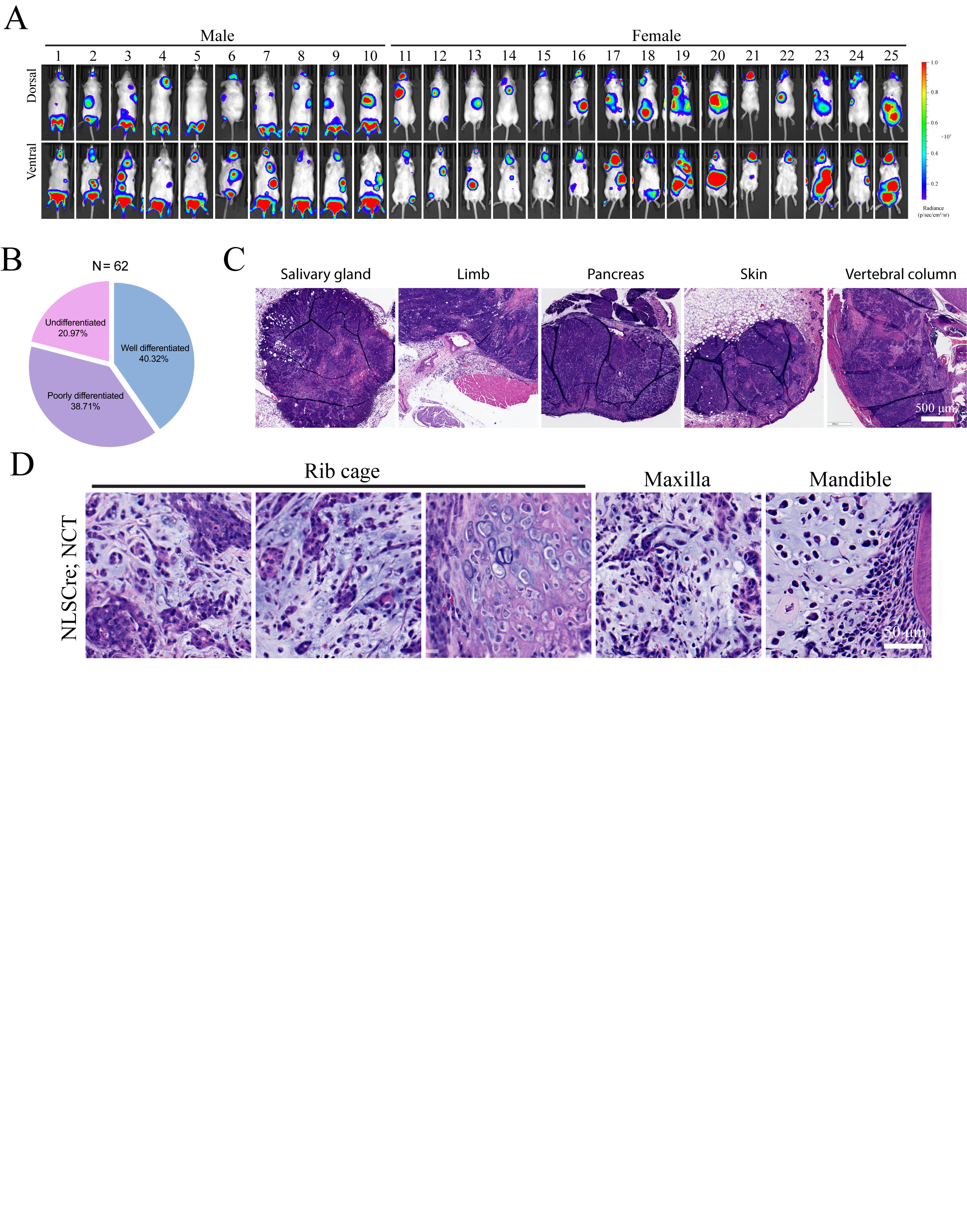

### Supplementary Figure 6

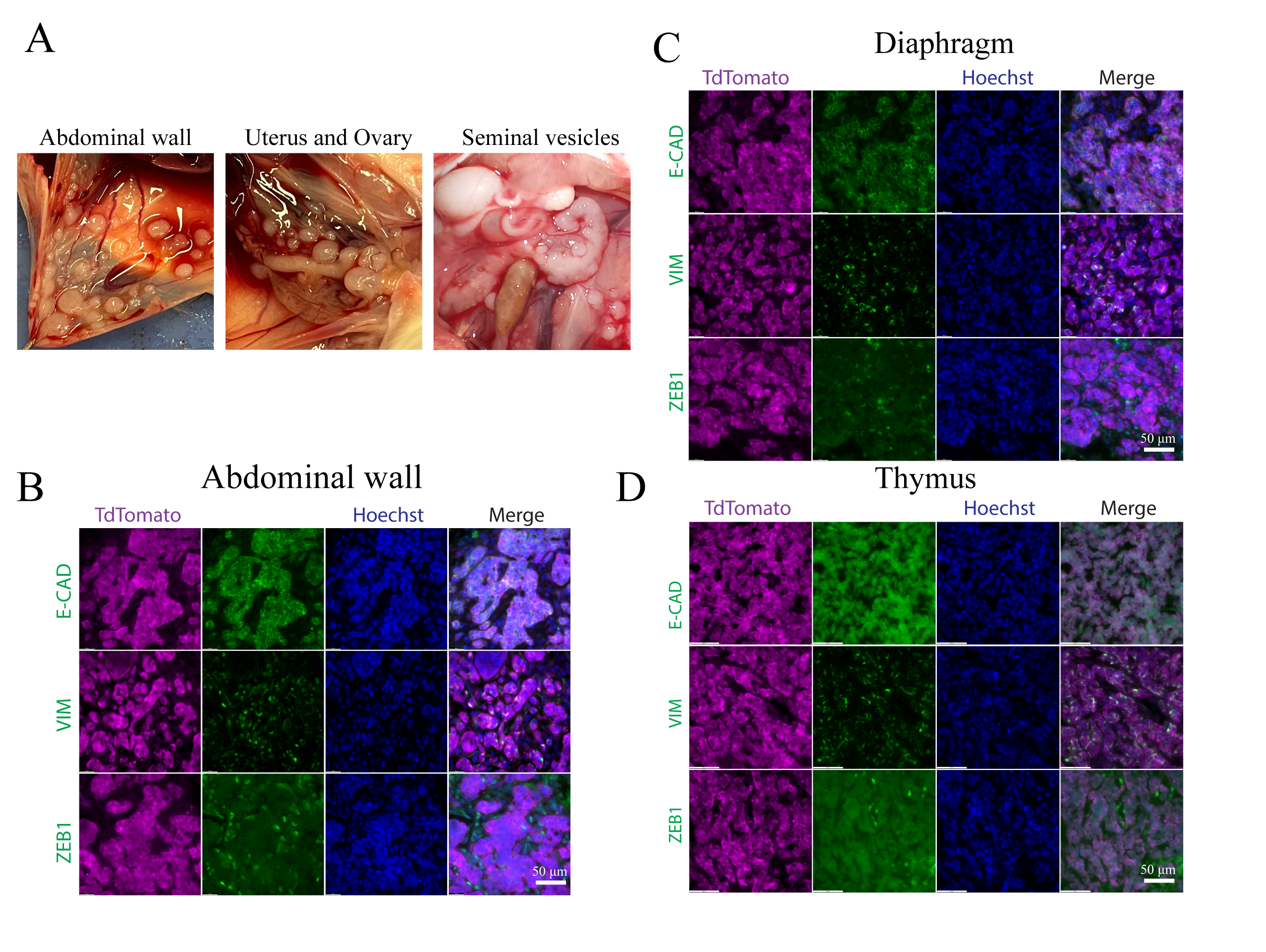
